# Pausing after clap reduces power required to fling wings apart at low Reynolds number

**DOI:** 10.1101/2020.10.26.356386

**Authors:** Vishwa T. Kasoju, Arvind Santhanakrishnan

## Abstract

The smallest flying insects such as thrips (body length < 2 mm) are challenged with needing to move in air at chord-based Reynolds number (*Re_c_*) on the order of 10. Pronounced viscous dissipation at such low *Re_c_* requires considerable energetic expenditure for tiny insects to stay aloft. Free-flying thrips flap their densely bristled wings at large stroke amplitudes, bringing both wings in close proximity of each other at the end of upstroke (‘clap’) and moving their wings apart at the start of downstroke (‘fling’). From high-speed videos of free-flying thrips, we observed that their forewings remain clapped for approximately 10% of the wingbeat cycle before start of fling. We sought to examine if there are aerodynamic advantages associated with pausing wing motion after clap and before fling at *Re_c_*=10. A dynamically scaled robotic clap-and-fling platform was used to measure lift and drag forces generated by physical models of non-bristled (solid) and bristled wing pairs for pause times ranging between 0% to 41% of the cycle. In both solid and bristled wings, varying pause time showed no effect on average force coefficients generated within each half-stroke. This was supported by nearly identical time-variation of circulation of the leading and trailing edge vortices for different pause times. At smaller pause times, bristled wings showed larger reduction of cycle-averaged drag coefficient as compared to that of solid wings. For a given wing design (solid or bristled), the ratio of cycle-averaged lift coefficient to cycle-averaged drag coefficient was unchanged across different pause times. We observed 13.5% drop in cycle-averaged power coefficient and 3% drop in cycle-averaged lift coefficient when moving from 0% pause to 9% pause duration. Our results suggest that pausing at the end of clap can be beneficial for reducing the power required to fling, with a small reduction in lift.

## 1 Introduction

Despite the roughly tenfold increase in wing length of a hawk moth compared to that of a fruit fly, the aerodynamic mechanisms underlying their free-flight are surprisingly similar. A vast number of studies examining flight aerodynamics of fruit flies and larger insects have identified the following mechanisms of lift generation: 1) delayed stall via the leading edge vortex (LEV) (Dickinson & Götz 1993, Ellington et al. 1996); 2) rotational lift (Dickinson et al. 1999, Sane & Dickinson 2002); 3) wing-wake interactions (Dickinson et al. 1999); and 4) wing-wing interaction during stroke reversal via clap-and-fling (Weis-Fogh 1973, Weis-Fogh 1975, Spedding & Maxworthy 1986). Far little is known about flight aerodynamics in entire families of miniature insects of body lengths ranging from 0.1 mm to 2 mm, such as thrips and several parasitoid wasps (e.g., *Trichogramma* spp. (Jalali et al. 2016) and fairyflies (Huber et al. 2008)). Miniature insects have been primarily examined by entomologists owing to their ecological and agricultural importance (Crespi et al. 1997, Terry 2001, Ullman et al. 2002, Whitfield et al. 2005, Jalali et al. 2016). From an engineering standpoint, studies of tiny insect flight can guide the development of biomimetic micro aerial vehicles (Liu et al. 2016).

Viscous dissipation of kinetic energy presents a significant constraint to the flight of tiny insects, where Reynolds number based on wing chord and tip velocity (*Re_c_*) is on the order of 1 to 10 (Santhanakrishnan et al. 2014, Jones et al. 2016, Santhanakrishnan et al. 2018). At such low *Re_c_*, these insects have to continually flap to stay aloft (Sane 2016). Multifold increase in drag coefficient has been reported for revolving (Santhanakrishnan et al. 2018) and translating (Miller & Peskin 2004) wings for *Re_c_* ≤32. At *Re_c_* ≥ 120 corresponding to flight of fruit flies and larger insects, a large LEV is formed at the start of a half-stroke and remains attached to the wing until the end of the halfstroke (Ellington et al. 1996, Birch et al. 2004). The trailing edge vortex (TEV) is detached from the wing and shed in the wake. The attached LEV delays stall and helps in lift generation (Dickinson et al. 1999, Ellington 1999). In contrast, both the LEV and TEV do not separate from a wing during linear translation (Miller & Peskin 2004) and revolution for *Re_c_* ≤ 32 (Santhanakrishnan et al. 2018). This LEV-TEV ‘vortical symmetry’ has been proposed to decrease lift in tiny insect flight (Miller & Peskin 2004), due to reduction in the time rate of change of the first moment of vorticity (Wu 1981).

Despite the above aerodynamic challenges, several studies have reported controlled flight of thrips over short distances (Terry 2001, Whitfield et al. 2005, Morse & Hoddle 2006, Rodriguez-Saona et al. 2010, Riley et al. 2011). Examining biomechanical adaptations used by tiny insects can help to understand how they are able to overcome fluid dynamic constraints. Two such key adaptations have been examined in several studies, including the presence of long bristles in their wings and obligatory use of wing-wing interaction in free-flight (clap-and-fling). Sunada et al. (2002) used dynamically scaled models undergoing translation and rotation and found little variations in forces between solid (non-bristled) and bristled wing designs. Force coefficients for the bristled wing model were found to be more compared to solid wing model, when using a reduced surface area to determine the coefficients of the bristled wing. Weihs & Barta (2008) and Davidi & Weihs (2012) found that a comb-like wing could comparatively generate forces similar to that of solid wings of same shape, while saving up to 90% of the wing weight. Recent studies (Lee & Kim 2017, Lee et al. 2018, Lee et al. 2020) have shown that a comb-like wing can provide aerodynamic benefit at small inter-bristle gaps, owing to the formation of diffused shear layers around the bristles that block flow from leaking through the gaps. However, most of these studies used a single bristled wing model and did not address wing-wing interaction used in free-flight of tiny insects (Lehmann et al. 2005).

Santhanakrishnan et al. (2014) performed 2D computational simulations of clap-and-fling at *Re_c_* corresponding to tiny insect flight. By approximating bristled wings as porous surfaces, this study found that bristled wings can provide substantial drag reduction when compared to solid wings during clap-and-fling. Jones et al. (2016) modeled wing bristles as 2D cylinder arrays and showed that bristled reduce the force required to fling the wings apart during wing-wing interaction. In our recent study (Kasoju et al. 2018), we experimentally examined the inter-bristle flow during clap-and-fling for bristled wing models with varying inter-bristle gap. When compared to a solid wing model, we found that bristled wings provide aerodynamic benefit through larger drag reduction and disproportionally lower lift reduction. Ford et al. (2019) found that thrips wings show a preference for smaller membrane area compared to the total wing area, and that wings with smaller membrane areas provide larger aerodynamic benefit during clap-and-fling at *Re_c_* corresponding to tiny insect flight. Collectively, these studies show that combining biomechanical adaptations in wing kinematics (clap-and-fling) and wing morphology (bristles) can provide aerodynamic benefit to flapping flight at the scale of the smallest insects.

In addition to the obligatory use of clap-and-fling, tiny insects have been observed to use a shorter upstroke duration and a longer downstroke duration (Santhanakrishnan et al. 2014). Such an asymmetric reduction of upstroke duration can lower the time where loss of lift occurs, as most of the lift in flapping flight of insects is generated during the downstroke (Sane 2003). Ellington (1975) observed that the tiny chalcid wasp *Encarsia Formosa* paused wing motion at the end of upstroke (clap) for about 10% of total cycle time (taken here as the sum of upstroke and downstroke time). He proposed that pausing at the end of clap could potentially promote shedding and advection of vortices away from the wing before the start of fling, and reduce the mechanical energy required for fling. These hypotheses were not tested in his study, and the aerodynamic implications of pausing after clap are unknown. In this study, we experimentally examine force generation during clap-and-fling at *Re_c_* = 10 across varying pause duration following the clap phase. Our tests were conducted using a dynamically scaled robotic model outfitted with bristled wing and solid wing physical models (Kasoju et al. 2018, Ford et al. 2019). 2D particle image velocimetry (PIV) measurements were used to examine the evolution and dissipation of flow structures around the wings during the pause following the clap phase.

## 2 Materials and methods

### 2.1. Free-flight recordings of thrips

Thrips were collected in Chapel Hill, NC, USA, during early June, 2017 from daylilies, gardenia and Azaleas flowers. The flowers with insects were then brought to recording arena and filmed within few hours of their collection. We used a procedure similar to that described in the study by Santhanakrishnan et al. (2014) for filming free take-off flight. A pipette tip was placed on top of an insect to allow it to crawl inside the tube. A single high-speed camera (Phantom v7.1, Vision Research, Wayne, NJ, USA) was used for filming. The camera was fitted with a 55 mm micro-Nikkor lens, a Nikon PB-5 bellows with variable extension, and a 27.5 mm extension tube. The pipette tip with thrips was placed upside-down in the camera field of view, and we waited for the thrips to crawl out of the tube and take-off from the tip. The field of view was illuminated using a red light emitting diode (LED) array. A white diffuser placed in front of the camera aperture, with the pipette tip located in between camera and the LED array. Free take-off flight of thrips were filmed at different frame rates with a shutter duration ranging between 15–30 *μ**s*** (Table 1).

**Table 1.** Pause duration analyzed between end of upstroke and start of downstroke for several high speed video recordings.

| Trial | Recording rate [fps] | Cycle time [ms] | Pause time [% cycle] |
| --- | --- | --- | --- |
| 1 | 3000 | 4.67 | 14.3 |
| 2 | 4000 | 5.25 | 9.5 |
| 3 | 4000 | 5.5 | 9.1 |
| 4 | 4000 | 4.75 | 10.5 |
| 5 | 4700 | 4.04 | 10.5 |

Five high-speed video recordings (representative snapshots shown in Figure 1) were digitized and analysed in ImageJ (National Institutes of Health, Bethesda, MD, USA) for calculating the pause time between the end of upstroke (clap) and start of downstroke (fling), and the results are provided in Table 1. The five raw videos that were used for analysis are provided as supplementary material (Movies S1-S5). The average pause time from the five recordings was calculated to be 11±2% of the total cycle time. This calculated pause time was close to that of *E. formosa* (10% of cycle time) reported by Ellington (1975).

**Figure 1.**
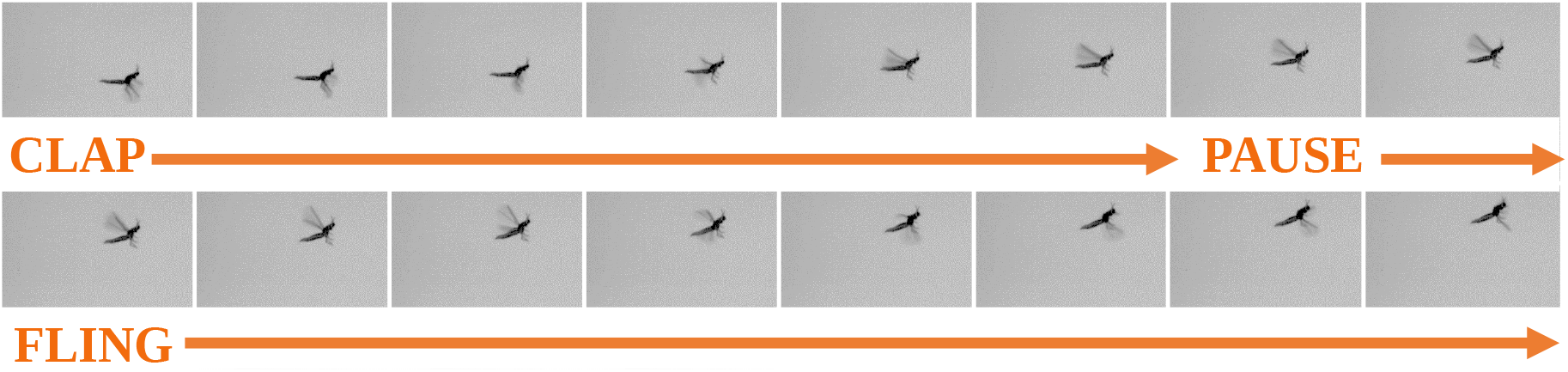
Successive snapshots of thrips in free takeoff flight. At the end of upstroke, both fore wings were brought in close proximity of each other (‘clap’). The wings paused for approximately 10% of flapping cycle before the start of downstroke (‘fling’). See Table 1 for more information.

### 2.2. Test facility

The dynamically scaled robotic wing platform used in this study has been used before (Kasoju et al. 2018, Ford et al. 2019) and is described briefly here. The robotic platform consists of four 2-phase hybrid stepper motors with integrated encoders (ST234E, National Instruments Corporation, Austin, TX, USA) mounted on an acrylic tank with 0.51 m x 0.51 m cross-section, and 0.41 m in height. These motors were used to prescribe the motion of 2 physical wing models. The four stepper motors were controlled by a multi-axis controller (PCI-7350, National Instruments Corporation, Austin, TX, USA) via custom programs written in LabVIEW software (National Instruments Corporation, Austin, TX, USA). Two stepper motors were dedicated to each wing to perform rotation and translational motion with help of bevel gear pairs and rack and pinion mechanism, respectively.

### 2.3. Physical models

A bristled wing of membrane width 7 mm, with symmetric bristle lengths on either side of a membrane (Figure 2A) was laser cut from an optically clear acrylic sheet of thickness 3.175 mm. The bristles were cut to required length from 304 stainless steel wires of uniform diameter (D)=0.3048 mm and were glued on top of the acrylic membrane with inter-bristle spacing (G) to bristle diameter (D) ratio (i.e., G/D) of 5 (Figure 2A). Also, an equivalent solid wing pair with the same chord and span lengths as the bristled wing model was laser cut from a 3.175 mm thick acrylic sheet. Each wing of the wing pair being tested was attached to custom made aluminum L-brackets and completely immersed inside the acrylic tank (described above under test facility) using 6.35 mm diameter stainless steel D-shafts.

**Figure 2.**
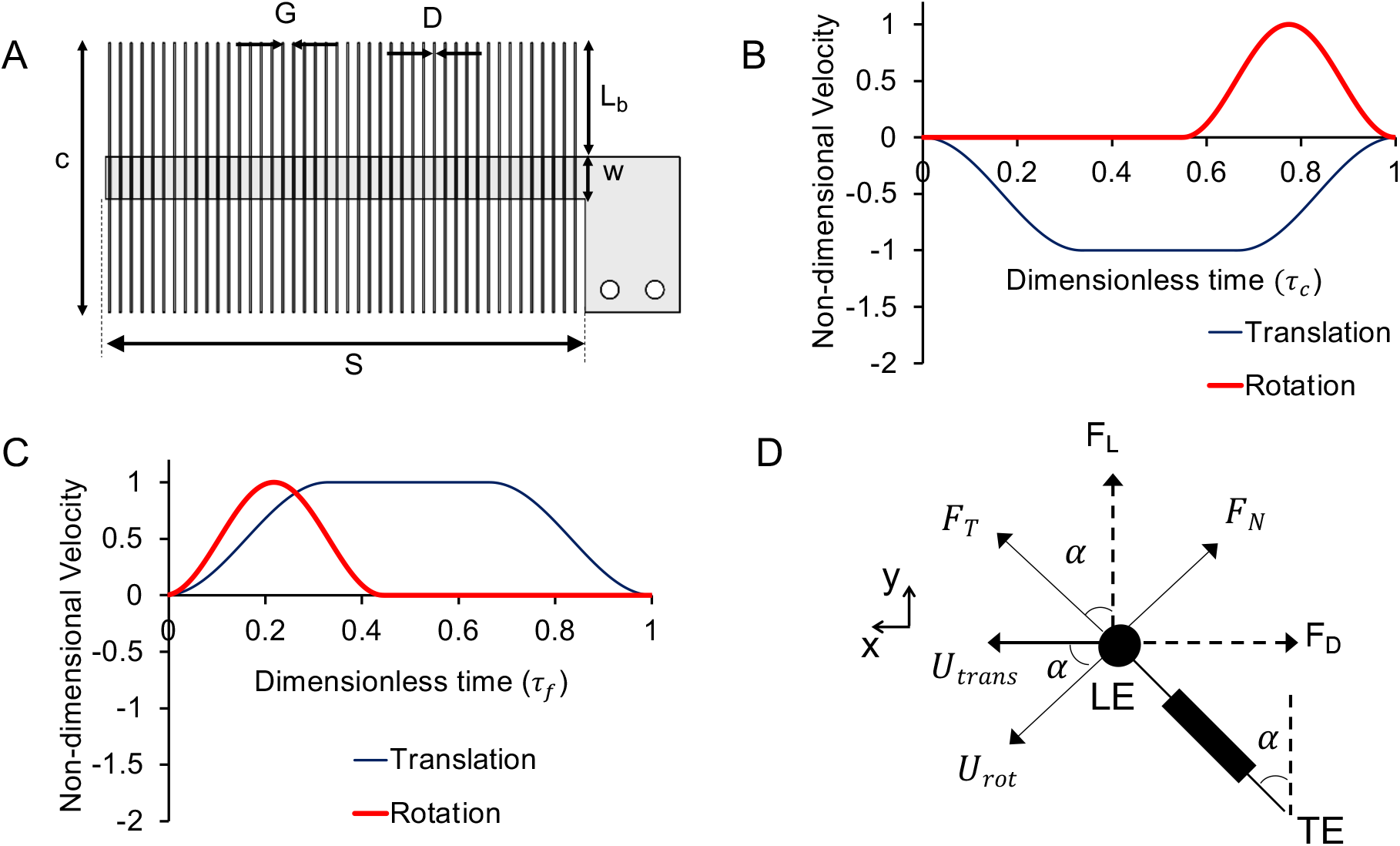
(A) Bristled wing model of chord length (c)=45 mm, wing span (S)=81 mm, inter-bristle spacing (G)=1.83 mm, bristle diameter(D)= 0.3048 mm, length of bristle (L_b_)=19 mm and membrane width (w)=7 mm. A solid wing model (without bristles) with the same chord (c) and span (S) lengths as that of the bristled wing was also tested. (B) and (C) show the time-varying motion profile prescribed for motion of a single wing during clap and fling, respectively, based on a previous study by Miller and Peskin (2005). The thin line indicates the wing translational motion while the thick line represents the wing rotation. (D) The sectional view of a bristled wing model (referred here as “chordwise view”) with directions of measured tangential *(F_T_*) and normal forces *(F_N_)* experienced during rotation by angle α. Lift (F_L_) and drag (*F_D_*) forces were measured by taking components of F_T_ and *F_N_* in the vertical and horizontal directions, respectively. τ_c_=dimensionless clap time; *τ_f_*=dimensionless fling time; LE=leading edge; TE=trailing edge; *U_trans_*=translational velocity at wing tip; *U_rot_*=rotational velocity at wing tip; x,y are global horizontal and vertical coordinate axes.

### 2.4. Wing kinematics

A modified version of 2D clap and fling kinematics that was initially developed by Miller & Peskin (2005) was prescribed for wing motion in the robotic model (Figure 2C–Figure 2D). The wings were made to rotate and translate simultaneously with 100% overlap prescribed between rotation and translation during both clap and fling phases. Wing rotation during clap was adjusted such that rotation ended exactly when the wings stopped translating, as shown in Figure 2C. During fling, wings were made to start rotation and translation at the same time, as shown in Figure 2D. Arora et al. (2014) previously examined the effects of varying the percentage of overlap between rotation and translation on forces generation, and reported increase in force coefficients with increasing overlap) during clap and fling. This was the rationale for choosing maximum possible overlap for both clap and fling in this study. Figure 2C and Figure 2D show prescribed non-dimensional velocities as a function of dimensionless time (*τ_c_,τ_f_*) during clap and fling, respectively. The dimensionless times for each phase (clap or fling) are indicated as the ratio of instantaneous time to total time of a specific phase (clap or fling). Note that the kinematics presented here are for a single wing performing clap and fling. The kinematics for the other wing were identical but in opposite directions. The inter-wing spacing between the wings was set to 10% of chord, which is similar to those observed in free flight recordings of thrips (Santhanakrishnan et al. 2014).

### 2.5. Test conditions

Force measurements and flow visualization were conducted for 5 pause times (0%, 9%, 17%, 23%, 41% of the entire cycle time). The total cycle time is calculated as sum of clap time, pause time and fling time, in units of milliseconds (ms). *Re_c_* = 10 was maintained as a constant across all test conditions, where *Re_c_* was based on steady translational velocity (U_st_) of the wing and chord length (c). The acrylic tank described in test facility above was filled with 99% glycerin solution to obtain *Re_c_*=10. The kinematic viscosity of the 99% glycerin solution used in this study was measured using a Cannon-Feske routine viscometer (size 400, Cannon Instrument Company, State College, PA, USA) to be 706×10^−6^m^2^/s. The density of the 99% glycerin solution was measured to be 1260 kg/m^3^. The corresponding Reynolds number based on bristle diameter (*Re_b_*) was 0.067 (based on steady translational velocity), which is within the biologically relevant range for tiny insect flight (Jones et al. 2016).

### 2.6. Force measurements

Forces on the wings were measured using uniaxial strain gauges bonded to the L-brackets (wing mount) and were designed to measure forces in perpendicular and parallel directions to the wing. These forces were then resolved along the global horizontal and vertical axes as drag and lift forces, respectively. Separate L-brackets were used for measuring lift and drag as described in a previous study (Kasoju et al. 2018). A data acquisition board (NI USB-6210, National Instruments Corporation, Austin, TX, USA) was used to acquire the strain gauge voltage data and angular position of the wings at a sample rate of 10 kHz throughout the entire cycle (includes clap time, pause time and fling time). We used the same processing procedures as used in Kasoju et al. (2018) and Ford et al. (2019). The raw data was filtered in MATLAB (The Mathworks Inc., Natick, MA, USA) using a third order low-pass Butterworth filter with a cutoff frequency of 24 Hz. The lift and drag brackets were calibrated manually and the calibrations were applied to the filtered voltage data. The forces were then resolved along global horizontal (drag force) and vertical (lift force) directions.

Dimensionless lift (*C_L_*) and drag (*C_D_*) coefficients were calculated as:

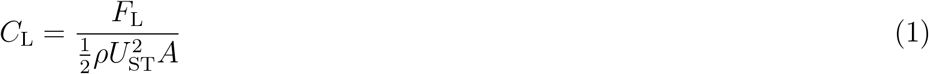

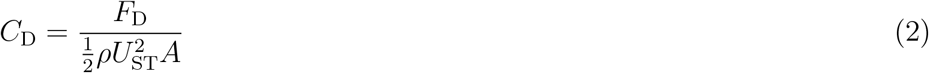

where *F_L_* and *F_D_* are the lift and drag forces measured along horizontal and vertical directions, respectively (in Newtons), *U_ST_* represents steady translational velocity, *ρ* is density of the fluid medium and *A* represents the effective wing surface area (4.05×10^−3^m^2^) for both the solid and bristled wing. The reason for using effective surface area for the bristled wing, as opposed to a reduced surface area (excluding gaps between the bristles), is because a reduced surface area implicitly assumes that flow through the bristles is mostly identical to the ideal/inviscid case without allowing the possibility that flow can incompletely leak through the gaps between the bristles on account of viscous interactions (Kasoju et al. 2018). Standard deviations were calculated across 30 consecutive cycles for *C_L_* and *C_D_*, and the force coefficients were averaged across all cycles. In addition, time-averaged force coefficients 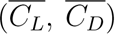 were calculated in clap, fling and throughout the entire cycle, with standard deviations and averages reported across all 30 cycles. We note that forces were only recorded on a single wing of a wing pair, with the assumption that force generation by other wing was symmetrical and equal in magnitude because the motion was symmetrical.

Similar to force coefficients, the power coefficient (*C_P_*) was calculated using the equation:

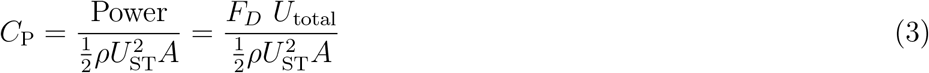

where *U*_total_ = *U*_trans_+*U*_rot_ cos α. *U*_trans_ and *U*_rot_ represents the wing tip velocity during translation and rotation, respectively, and α represents the wing rotation angle shown in Figure 2D.

### 2.7. Flow visualization

2D time-resolved PIV (2D TR-PIV) were conducted to visualize and measure the flow generated during the clap phase, pause duration and fling phase by the solid and bristled wing pairs along chordwise direction at the mid-span location (wings in chordwise view similar to Figure 2D). 2D TR-PIV measurements were acquired for both wing models (solid and bristled) at all test conditions (0%, 9%, 17%, 23%, 41% pause time). A single cavity Nd:YLF laser (Photonics Industries International, Inc., Bohemia, NY, USA) was used for illumination that provided a 0.5 mm diameter beam of 527 nm in wavelength. A thin laser sheet (thickness ≈3-5 mm) was generated from the beam using a cylindrical lens of 10 mm focal length. A high-speed complementary metal-oxide-semiconductor (CMOS) camera with a spatial resolution of 1280×800 pixels, maximum frame rate of 1630 frames/s, and pixel size of 20×20 microns (Phantom Miro 110, Vision Research Inc., Wayne, NJ, USA) was used for acquiring raw TR-PIV images. This camera was fitted with a 60 mm constant focal length lens (Nikon Micro Nikkor, Nikon Corporation, Tokyo, Japan). Hollow glass spheres of 10-micron diameter (110P8, LaVision GmbH, Gottingen, Germany) were used as seeding particles (Kasoju et al. 2018, Ford et al. 2019). 100 evenly spaced images were acquired at a recording rate of 90 Hz during the clap and during the fling. The raw images were processed using DaVis 8.3.0 software (LaVision GmbH, Göttingen, Germany) using the following crosscorrelation settings: one pass with an interrogation window of size 64×64 pixels and two subsequent passes with interrogation window of size 32×32 pixels, each with 50% overlap. The processed 2D TR-PIV images were phase-averaged over 5 non-consecutive cycles. Following phase-averaging, 2D velocity vector fields were exported for calculating circulation (Γ) of the LEV and the TEV on a single wing of the imaged wing pair. Γ was calculated for 12 equally-spaced time points (every 5% of cycle time) during clap and fling separately using an in-house MATLAB script using Stokes’ equation:

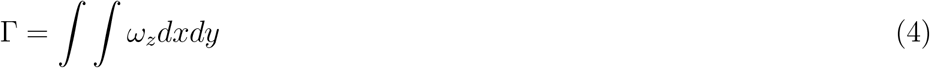

where *ω_z_* represents the out-of-plane (z-component) of vorticity at the leading or trailing edges calculated from exported velocity vector fields (similar to Ford et al. (2019)) and *dxdy* represents the vorticity region for either the LEV or the TEV. We used a minimum cutoff of 10% of the maximum overall vorticity (*ω_z_*) at the leading and trailing edges across all the time points tested for clap and fling phases separately. Circulation (Γ) on a wing was calculated by drawing a box around LEV or TEV separately, and integrating the vorticity of the closed contour using equation 4. Γ was determined for the right-hand side wing only, with the assumption that circulation for the left wing will be equivalent in magnitude but oppositely signed. Note that the left wing motion is symmetric to right wing making our assumption justifiable.

2D phase-locked PIV (2D PL-PIV) measurements were acquired for wing models along a spanwise plane (similar to 2D PL-PIV in Kasoju et al. (2018)) located at 50% of bristle length (L_b_), measured from the membrane to the leading edge of the wing (Figure 2A). A double-pulsed, single-cavity Nd:YAG laser (Gemini 200-15, New Wave Research, Fremont, CA) with wavelength of 532 nm, maximum repetition rate of 15 Hz, and pulse width in the range of 3-5 ns was used for illumination in the PL-PIV measurements. A 10 mm focal length cylindrical lens was used to generate a thin laser sheet (thickness ≈3-5 mm). Raw PL-PIV images were acquired using a scientific CMOS (sCMOS) camera, with a maximum spatial resolution of 2600×2200 pixels at a frame rate of 50 frames/s, and a maximum pixel size of 6.5×6.5 microns (LaVision Inc., Ypsilanti, MI, USA). The 60 mm lens used in TR-PIV measurements was also used for PL-PIV measurements, and the camera was focused on seeding particles (hollow glass spheres, 10-micron diameter) along the laser plane (Kasoju et al. 2018). Raw image pairs were acquired at 7 time points in fling at equally spaced time steps of 12.5% of stroke times (*τ_f_*). The laser pulse separation between the images of an image pair were estimated based on 6-8 pixels of particle movement from one image to other image. For each wing model tested at *Re_c_* of 10, 5 image pairs were acquired at each time point fling cycle from 5 continuous cycles of clap and fling. These raw image pairs were processed using DaVis 8.3.0 software (LaVision GmbH, Göttingen, Germany) and then averaged for each time point. The post-processing parameters for 2D PL-PIV measurements were the same as those described earlier in 2D TR-PIV. The averaged processed images were exported to quantify the amount of fluid leaked through the bristles along the wing span. (Cheer & Koehl 1987) estimated the amount of fluid leaking through a pair of cylinders using a non-dimensional index called leakiness (*L_e_*). Leakiness (*L_e_*) is defined as the ratio of the volumetric flow rate of fluid that is leaked through the inter-bristle gaps in the direction opposite to wing motion under viscous (realistic) conditions to the volumetric flow rate for inviscid conditions, and is given by the equation:

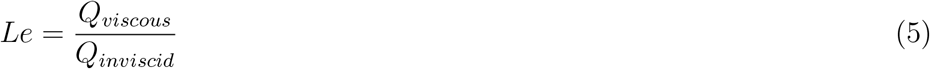

where *Q_viscous_* represents the volumetric flow rate leaked through the bristles under viscous condition calculated from the 2D PL-PIV measurements alog wing span, *Q_inviscid_* represents the volumetric flow rate leaked through the bristles under no viscous resistance (inviscid flow) calculated based on the assumption that under no viscous resistance, all the flow leaks through the inter-bristle gap (Kasoju et al. 2018).

## 3 Results

### 3.1. Force generation

During clap, the trend for lift (*C_L_*) and drag (*C_D_*) coefficients (Figure 3) are consistent with previously published data on solid and bristled wings (Santhanakrishnan et al. 2014, Kasoju et al. 2018, Ford et al. 2019). Starting from rest, the two wing-pairs were made to rotate and translate towards each other showing an increase in force coefficients in the initial acceleration phase. This is followed by constant velocity wing translation (τ_c_=0.35-0.7), where both solid and bristled wing were found to show little variation in force generation (*C_D_* and *C_L_*) in time. During the end of clap phase (*τ_c_*= 0.7-1), we observed the drag coefficient (*C_D_*) to vary significantly in time for a solid wing (Figure 3A) compared to bristled wing (Figure 3B). This was presumably due to wing-wing interaction, as the wings approach close to each other at the end of clap phase. However, lift coefficients (*C_L_*) for both solid and bristled wings (Figure 3C, Figure 3D) were found to drop during the end of clap phase. Interestingly, changing the pause time after clap phase showed no variation in force generation, suggesting that the pause time have no prior affects on force generation.

**Figure 3.**
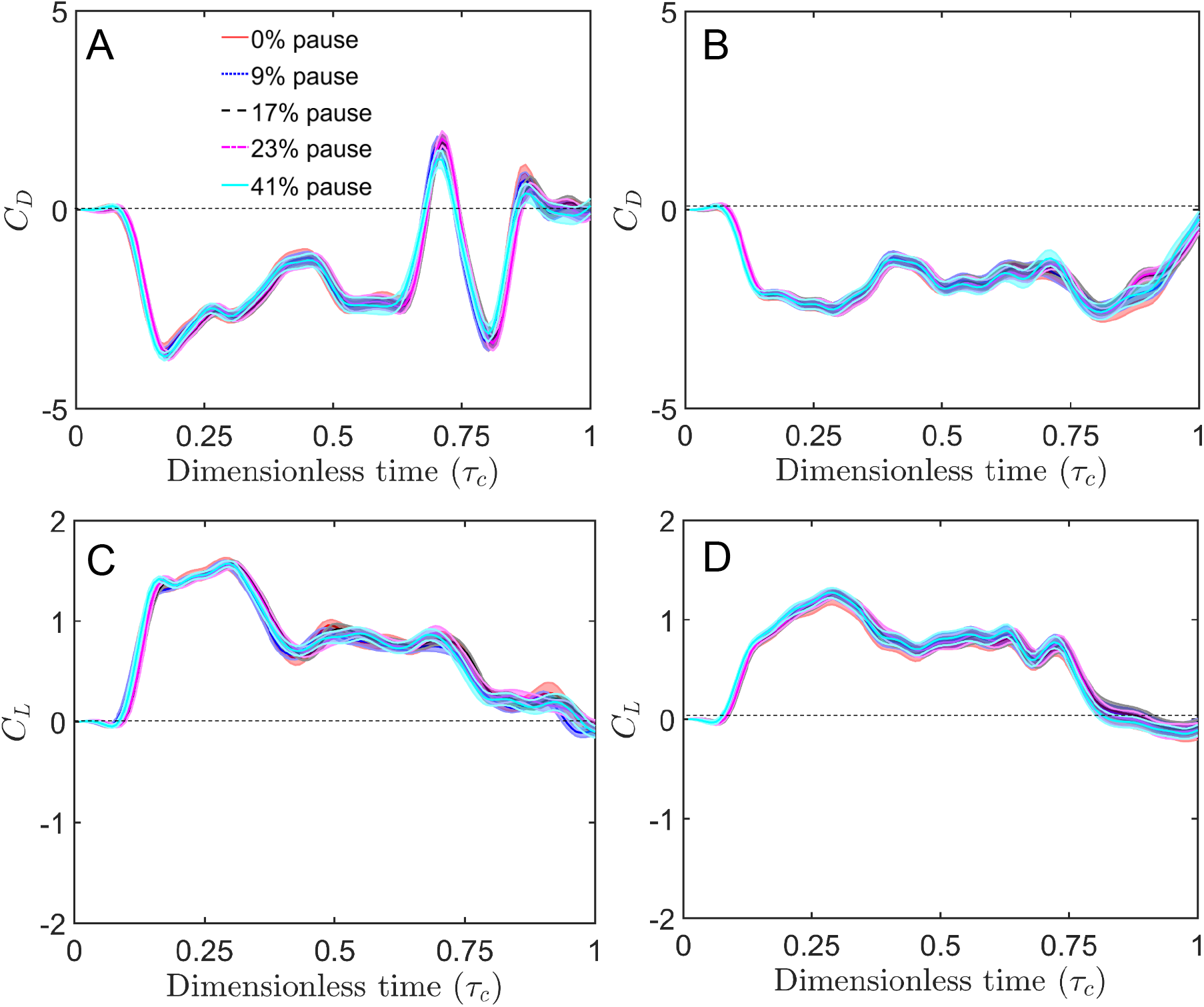
Force coefficients during clap at *Re_c_* =10 with shading around each curve representing range of ±1 standard deviation for that particular data (across 30 cycles). (A) and (C) show the drag coefficient (*C_D_*) and lift coefficient (*C_D_*), respectively, during clap *(τ_c_)* for solid wing model for various pause times before the start of fling. (B) and (D) show the drag coefficient (*C_D_*) and lift coefficient (*C_L_*) respectively during clap (τ_c_) for bristled wing model for various pause times.

Drag coefficient (*C_D_*) was found to peak during early stages of fling for both solid and bristled wings (Figure 4A, Figure 4B), where the wings were accelerating when performing rotation and translation. This tremendous increase in drag coefficient during early stages of fling was presumably due to wing-wing interaction. Interestingly, drag coefficients were found to drop for the rest of the fling phase, when the wings moved farther apart. This clearly indicates the influence of wing-wing interaction on drag coefficient, and was also observed in several previous studies (Miller & Peskin 2004, Miller & Peskin 2005, Arora et al. 2014, Santhanakrishnan et al. 2014, Jones et al. 2016, Kasoju et al. 2018, Ford et al. 2019). Peak *C_D_* for a bristled wing during fling was significantly lower compared to a solid wing for any test condition tested in this study (Figure 4A, Figure 4B). Increasing the percentage of pause time before the start of fling showed little to no variation in *C_D_*. This suggests that pause time has no significant effect on drag force generation in fling. Similar to *C_D_*, lift coefficient (*C_L_*) for both solid and bristled wings were found to peak in the early stages of fling suggesting the influence of wing-wing interaction. For the rest of the fling phase, *C_L_* was found to mostly remain constant and then drop resembling the constant velocity translation and deceleration of the wing, respectively. Similar to *C_D_*, influence of changes in pause time are minimal for *C_L_* during fling.

**Figure 4.**
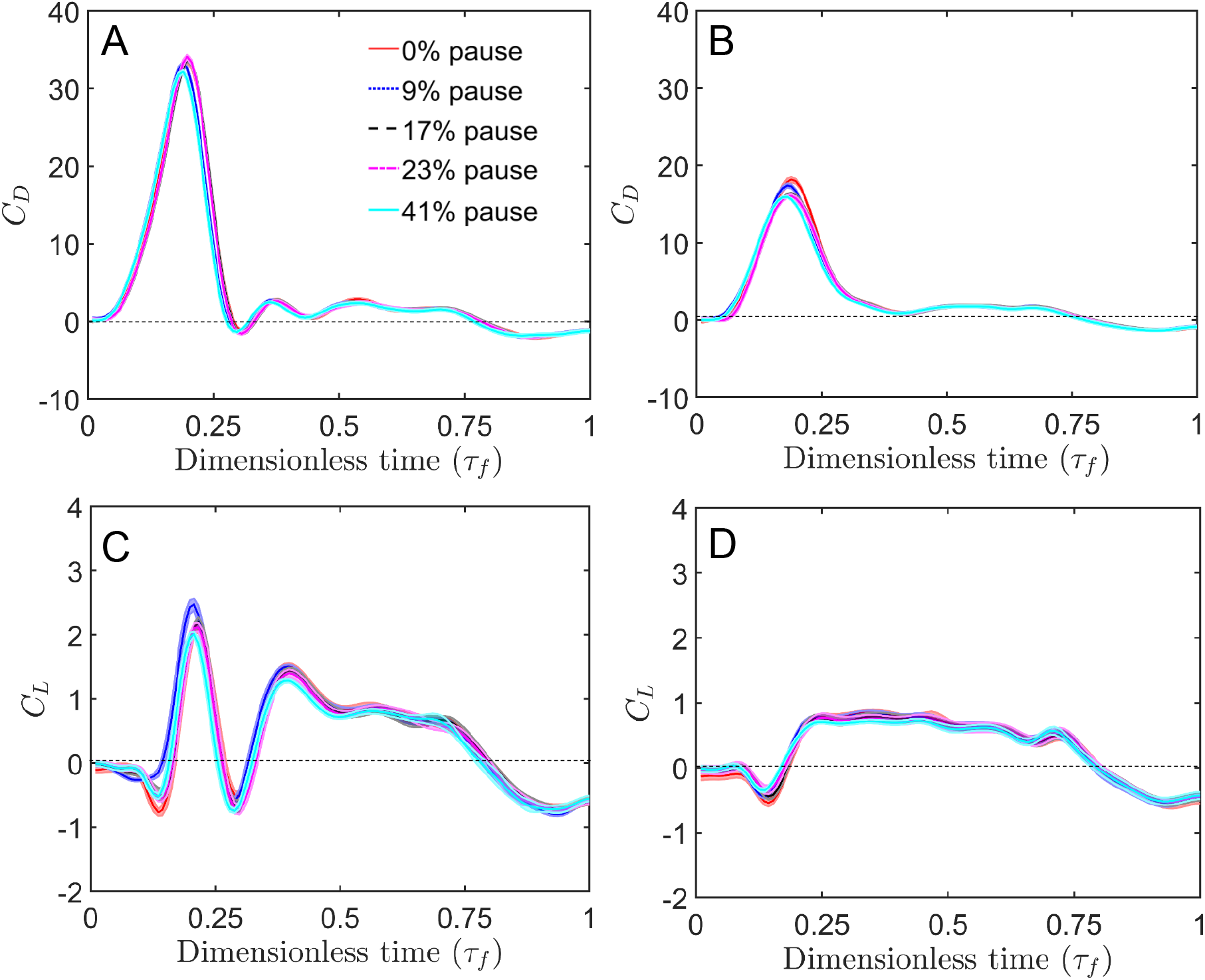
Force coefficients during fling at *Re_c_*=10 with shading around each curve representing range of ±1 standard deviation for that particular data (across 30 cycles). (A) and (C) show the drag coefficient (*C_D_*) and lift coefficient (*C_L_*), respectively, during fling (*τ_f_*) for solid wing model for various pause times before the start of fling. (B) and (D) show the drag coefficient (*C_D_*) and lift coefficient (*C_L_*), respectively, during fling (*τ_f_*) for bristled wing model.

To obtain an overall understanding of the changes in force coefficients (*C_D_* and *C_L_*) with pause time in between clap and fling phase, we examined average force coefficients (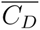 and 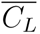) during clap and fling separately (Figure 5). Changes in pause time showed no influence on average force coefficients (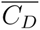 and 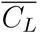) during clap for both solid and bristled wings (Figure 5A, Figure 5B). Interestingly, the values of 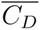 and 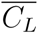 for both solid and bristled wings were almost similar in clap. During fling (Figure 5C, Figure 5D), changes in pause time also showed little to no influence on average force coefficients (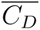 and 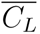). In contrast to clap, 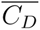during fling for solid wing was more compared to a bristled wing across all pause times. However, 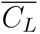was nearly the same for both solid and bristled wing at all pause times during fling.

**Figure 5.**
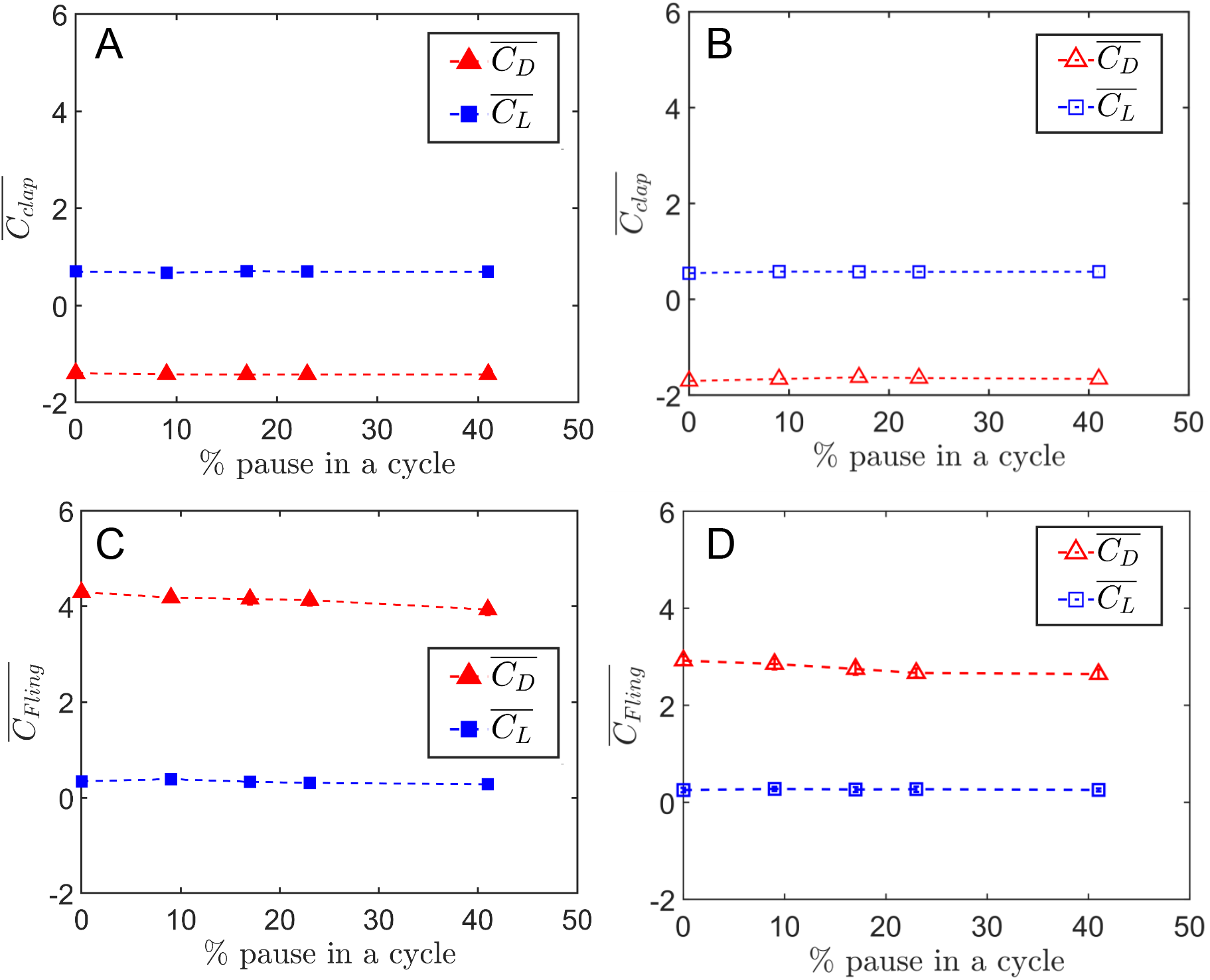
Average force coefficients during clap and fling at *Re_c_* = 10 are presented separately with error bars representing ±1 standard deviation for that particular data (across 30 cycles). (A) and (B) show the average drag coefficient (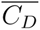) and average lift coefficient (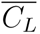) for varying pause times during clap for solid and bristled wing model, respectively. (C) and (D) show 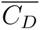 and 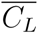 for varying pause times during fling for solid and bristled wing model, respectively.

### 3.2. Chordwise flow structures

Average force coefficients (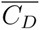 and 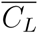) showed no variation between solid and bristled wing models during clap, and the flow structures were also essentially similar when comparing the solid and bristled wing models (Figure 6). The flow structures around a solid wing during clap were similar to those observed in our previous studies (Kasoju et al. 2018) and are thus not shown here. However, we examined the strength of the flow structures by calculating LEV circulation (Γ_LEV_) and TEV circulation (Γ_TEV_) of the solid and bristled wing models during clap (Figure 7A, Figure 7B) and during fling (Figure 7C, Figure 7D). Both Γ_LEV_ and Γ_TEV_ followed the same trend in time during clap when comparing solid (Figure 7A) and bristled (Figure 7C) wings. In addition, the magnitude of both Γ_LEV_ and Γ_TEV_during clap for the bristled wing model were similar to that of the solid wing, and this similarity is in agreement with the observed force generation during clap. Changing the pause time showed no variation in Γ_*LEV*_ and Γ_TEV_ during clap.

**Figure 6.**
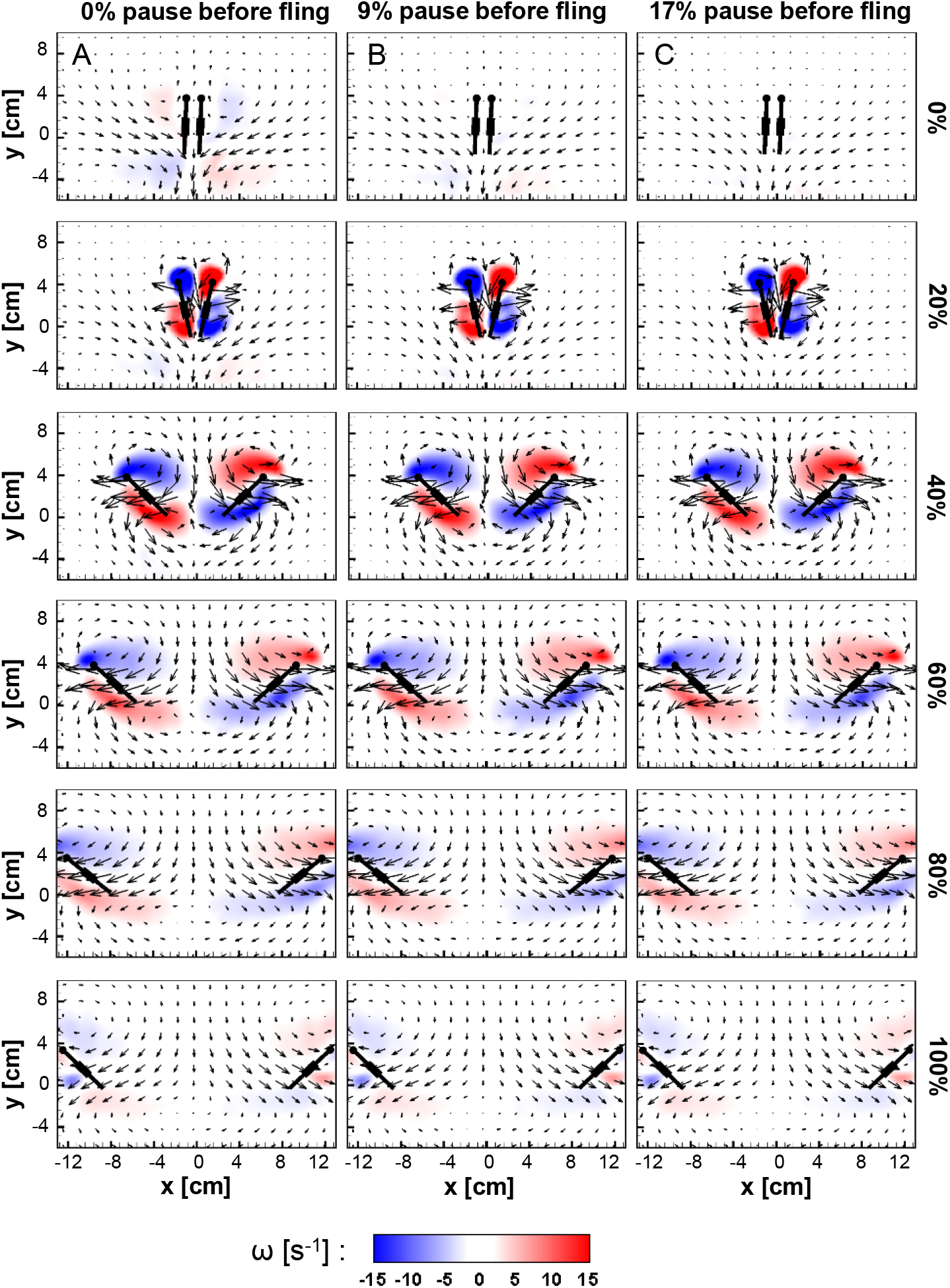
Velocity vector fields overlaid on out-of-plane z-vorticity (*ω_z_*) contours for bristled wing during fling at *Re_c_*=10 for various pause times: (A) 0%, (B) 9%, (C) 17% of cycle time. For each pause condition, 6 timepoints (0%, 20%, 40%, 60%, 8% and 100% of fling time) are shown along each column (increasing time from top to bottom). Red colour represents counterclockwise vorticity, while blue represents clockwise vorticity.

**Figure 7.**
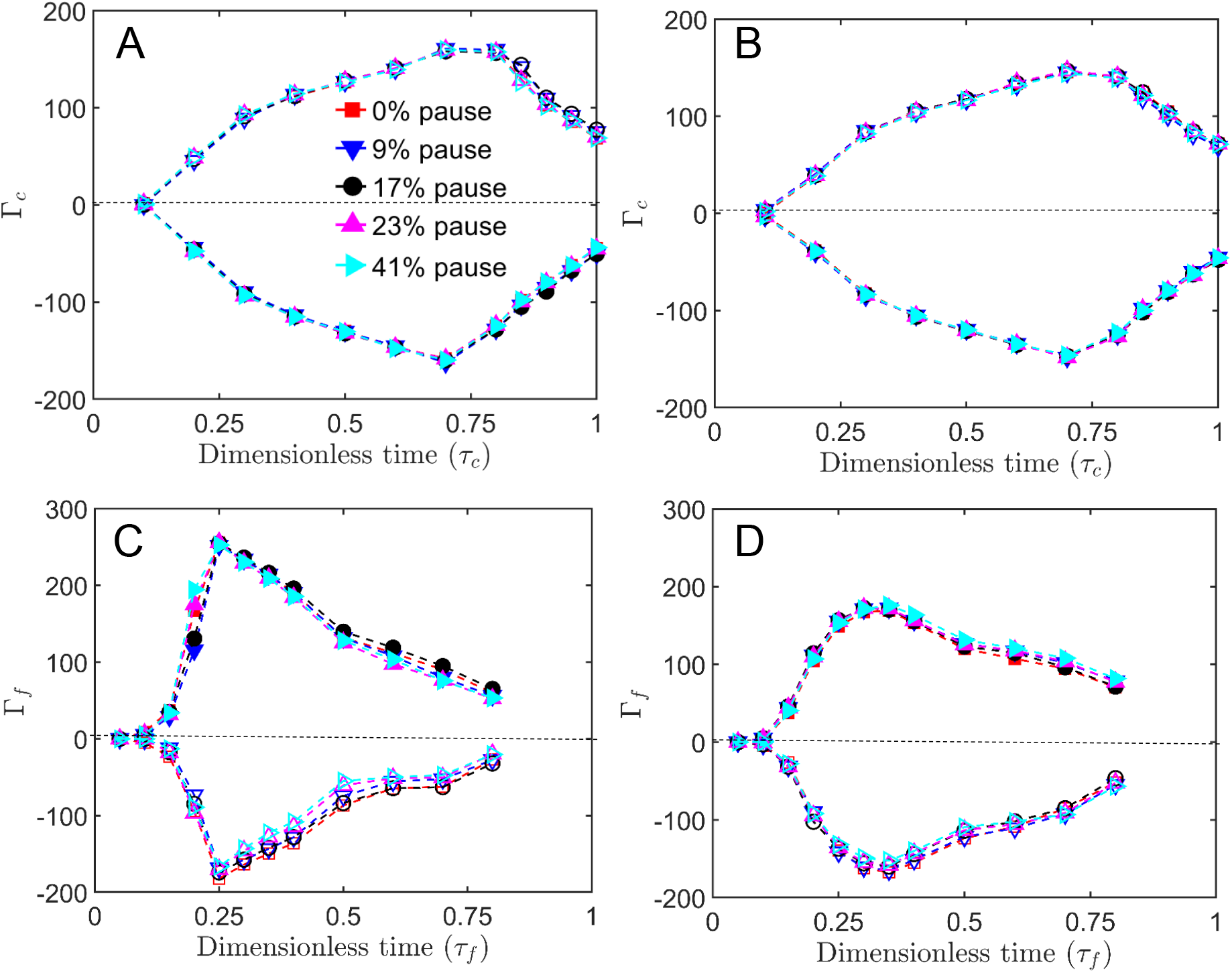
LEV AND TEV circulation as a function of dimensionless time. (A) and (C) show circulation on a solid wing during clap and fling, respectively. (B) and (D) show circulation on a bristled wing during clap and fling, respectively. Solid markers represent LEV circulation and hollow markers represent TEV circulation.

Similar to clap, flow around the wings showed little to no variation between solid and bristled wing models during fling. Both the LEV and the TEV were found to increase in strength during early stages of fling (Figure 6) and then found to decrease in strength with vorticity being diffused into the fluid medium surrounding the wing. For the solid wing model, circulation (Γ_LEV_ and Γ_TEV_) was found to steeply increase and decrease in time during early stages of fling (Figure 7C). While for the bristled wing, a steady increase and decrease in both Γ_*LEV*_ and Γ_*TEV*_ was observed (Figure 7D). With increasing the pause time before start of fling, no variation was observed throughout the fling time. Interestingly, just before the start of fling for 0% pause case, we observed the formation of a wake with low vorticity in the fluid medium surrounding the wings. This was most likely remnant of the wake generated from the clap phase that was just completed. However, the wake shed from clap was not found to affect the circulation around the wings during fling. With increasing pause time, we observed the wake remnant from clap to diminish before the start of fling.

## 4 Discussion

With the help of a robotic model, we measured forces and performed flow visualization on solid and bristled wing pairs during clap and fling under varying pause time at *Re_c_*=10. We find that including a pause time between clap and fling phases results in a negligible impact on average force coefficients (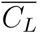 and 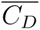) during clap (phase before pause) and during fling (phase after pause) for both solid and bristled wing models. This was also supported when examining LEV and TEV circulation, both of which showed no variation with changing pause time in either clap or fling. These results comprehensively suggest that including a pause between clap and fling phases has no influence on forces generated and flow structures formed during clap and fling at *Re_c_*=10.

High-speed video recordings of thrips showed that these insects pause for about 10% of the cycle time, which was in agreement with previous published results on flapping flight of the tiny wasp *E. formosa* (Ellington 1975). However, the influence of pause time on aerodynamic force generation have not been examined in previous studies of clap and fling at low *Re_c_* (Miller & Peskin 2004, Kasoju et al. 2018, Arora et al. 2014, Ford et al. 2019). In this study, a bristled wing model with total surface area equal to 33% of a geometrically similar solid wing area was tested. This drop in surface area of the wing should directly decrease the force generated by bristled wings(refer to equations (1), (2)). However, irrespective of pause time, the average force coefficients (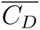 and 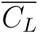) for a solid wing during clap were similar to that of bristled wing. We suspect that this similarity in forces between the solid and bristled wings is due to the blockage effect that is caused by the shear layers around the bristles at lower G/D ratios and has been described in previous studies (Lee & Kim 2017, Kasoju et al. 2018). This phenomenon causes the inter-bristle gap to be blocked due to shear layers formed around each bristle, thereby not allowing the fluid to pass through the gaps in between the bristles. This forces the fluid to move around the bristled wing (Davidi & Weihs 2012), thereby generating forces that are mostly equivalent to a solid wing model. Furthermore, this similarity between the solid and bristled wing was also evident in the circulation plots, where both Γ_*LEV*_ and Γ*_TEV_* were similar for both solid and bristled wing (Figure 7A,B).

Similar to the clap phase, average lift coefficient (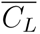) during fling was similar for both solid and bristled wing model irrespective of pause time. However, a drop in 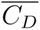 was observed for bristled wing compared to solid wing model. Further investigation of drag coefficient (*C_D_*) in time for a bristled wing model during fling showed that a 43% drop in peak drag coefficient relative to the solid wing model. Interestingly, investigating the flow through the bristles using 2D PL-PIV and characterizing the leakiness (*Le*), we found that the peak leakiness was about 40%, which is similar to drop in peak *C_D_* relative to the solid wing (Figure 8). We therefore conclude that leakiness is responsible for the observed drop in *C_D_* during fling.

**Figure 8.**
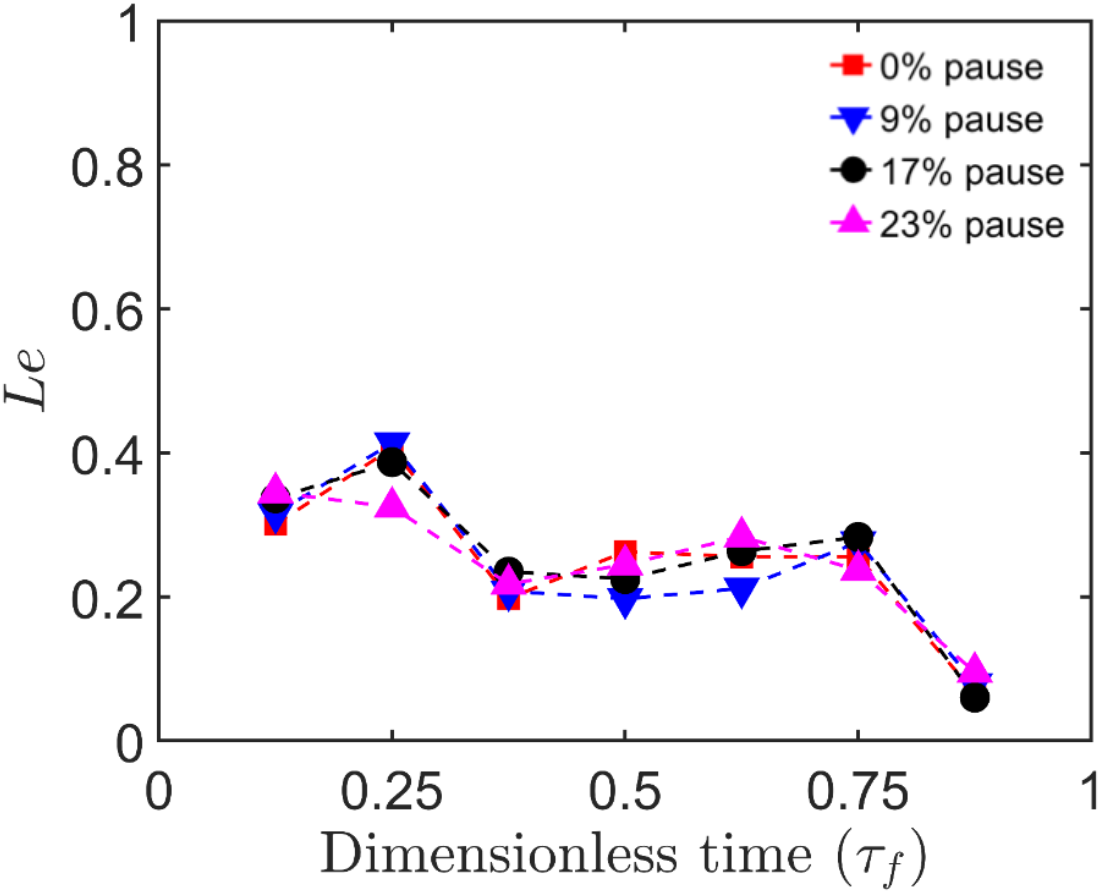
Leakiness (Le), representing non-dimensional flow reduction by the bristled wing, as a function of fling time (*τ_f_*) for various pause times.

Figure 9A and Figure 9B show 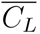 and 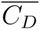 for the entire cycle (sum of clap time, pause time and fling time). Increase in pause time increases the entire cycle time. With increasing pause time, both 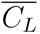 and 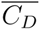 were found to decrease for solid wing model. Comparatively, when considering the standard deviations, increasing pause time resulted in marginal impact on 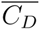 and 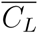 for larger pause times in the bristled wing model (Figure 9A, Figure 9B). As the % of pause time increases, both 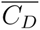 and 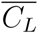 of the solid wing model was found to reach values close to that of bristled wing model. Significant drop in 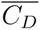for the bristled wing model (as compared to the solid wing) was realized at 0% pause time as compared to 41% pause time. Therefore, bristled wings can be beneficial for drag reduction at lower pause times. Further investigating the average lift to average drag ratio (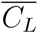/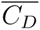) revealed that the aerodynamic performance of solid and bristled wings were essentially invariant with changing pause time (Figure 9C). These results potentially suggest that tiny insects that tend to pause their wing motion between clap and fling need not compromise their aerodynamic performance when doing so (assessed here using 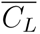/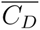).

**Figure 9.**
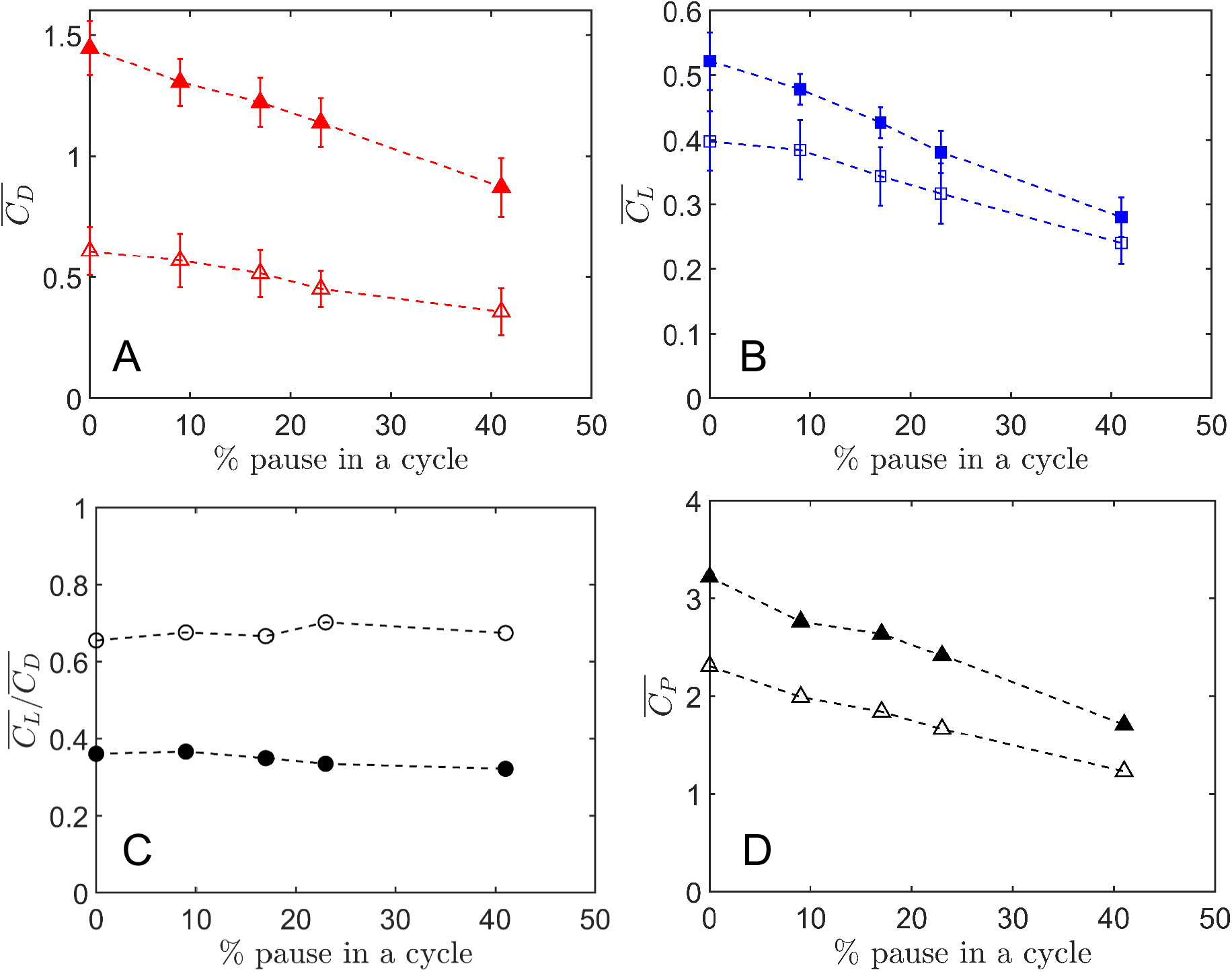
(A,B) Average force coefficients (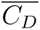, 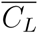), (C) average lift over average drag ratio (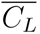/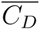), and (D) average power coefficient (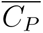) calculated over the entire cycle (clap time, pause time and fling time) across varying pause times. Solid markers represents solid wing model, hollow markers represents bristled wing model.

Ellington (1975) hypothesized that including a pause before the start of fling could help the insects in elastic storage of high mechanical energy that would be needed to fling the wings apart. Although elastic storage of energy in flight has not been examined for tiny insects such as thrips, a previous study by Dickinson and Lighton (Dickinson & Lighton 1995) presented clear evidence that fruit flies need elastic mechanisms for efficient flight. Another study by Alexander (1995) suggested that wing muscles of many insects can function as springs and store energy for reuse in the next flapping stroke. Another means to achieve efficient flight is to have effective muscle efficiency, which requires larger metabolic energy consumption. Whether or not thrips use elastic storage needs to be examined in future studies.

If we were to assume there is no elastic storage, tiny insects such as thrips require large muscle power to overcome severe viscous drag during the start of fling (downstroke) or during braking. A non-dimensional estimate of the power required to overcome the drag is presented in Figure 9D. Increasing the pause time results in a decrease of the average power coefficient (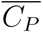), thus lowering the power required to perform clap and fling in both the solid and bristled wing models. However, increasing the pause time inevitably also results in decreasing lift. We observed a 13.5% drop in average power coefficient compared to a 3% drop in lift coefficient when increasing the pause time from 0% to 9% of the cycle. This provides evidence that pausing before fling can be beneficial for reducing the power required to overcome drag and perform clap and fling, albeit with a small compromise in lift. The pause time could serve as a “resting period” for insects, helping them to reduce their total power consumption.

## 5 Conclusions

This study showed that pause time between clap and fling has no influence on timevarying aerodynamic forces generated during clap and fling phases for both solid and bristled wing pairs at a Reynolds number of 10. However, considering the average force coefficients for the entire cycle (clap phase, pause time, fling phase), bristled wings were found to be beneficial in drag reduction at lower pause times. For 9% pause time between clap and fling, we observed a 13% drop in average power coefficient (*C_P_*) for a modest 3% reduction of the average lift coefficient. Collectively, our findings suggest that pausing before fling can help to reduce the power consumption in clap-and-fling, with a small compromise in lift.

## Supplementary Material

The raw high-speed recordings corresponding to 5 trials listed in Table 1 are available at: https://figshare.com/s/b5fa96f717fcd7561d1f. (Note: this is a private link for the review process and will be made public with a DOI if/when manuscript is accepted).

## Acknowledgments

The authors would like to thank the following collaborators at the University of North Carolina: Prof. Tyson L. Hedrick for his assistance with acquiring videos of free-flying thrips and providing us generous access to his lab facilities, and Prof. Laura A. Miller for her advice on collecting thrips. The authors would also like to thank the following students at Oklahoma State University: Mitchell P. Ford for pointing to Prof. C. P. Ellington’s observation on pausing after clap in *E. formosa*, and Truc T. Ngo for her assistance with digitizing thrips flight videos.

## Competing interests

The authors declare no financial or otherwise competing interests.

## Funding

This research was funded by the National Science Foundation (CBET 1512071 to AS).

## Notes

### Competing Interest Statement

The authors have declared no competing interest.

